# Motor, somatosensory, and executive cortical areas directly modulate firing activity in the auditory midbrain

**DOI:** 10.1101/2023.07.25.550491

**Authors:** Sarah E Gartside, Adrian Rees, Bas MJ Olthof

## Abstract

We have recently reported that the central nucleus of the inferior colliculus (the auditory midbrain) is innervated by glutamatergic pyramidal cells originating not only in auditory cortex (AC) but also in multiple ‘non-auditory’ regions of the cerebral cortex. Using optogenetics and electrical stimulation, we investigated the functional properties of these descending connections in vivo in anaesthetised rats. A retrograde virus encoding green fluorescent protein (GFP) and channelrhodopsin (ChR2) injected into the central nucleus of the inferior colliculus (ICC), labelled discrete groups of cells in multiple areas of the cerebral cortex. Light stimulation of AC and M1 caused local activation of cortical neurones and increased the firing rate of neurones in ICc indicating a direct excitatory input from AC and M1 to ICC. Electrical stimulation of M1, secondary motor, somatosensory and prefrontal cortical regions evoked short, fixed latency firing events in ICC as well as longer latency, longer duration increases in firing activity. The short latency events were singular spikes of consistent shape and size likely resulting from monosynaptic excitation of individual ICC units. The longer latency responses comprised multiple units and spikes occurred with significant temporal jitter suggesting polysynaptic activation of local circuits within the ICC. The probability of the monosynaptic event, the magnitude of the polysynaptic response, and the area of ICC affected were dependent on the stimulus current. Our data are consistent with cortical regions exerting an important excitatory direct and indirect regulation of ICc neurones.

**Significance statement:** We have recently described inputs from motor, somatosensory, and executive cortices to the inferior colliculus (IC, auditory midbrain). Here we provide functional evidence for such connections. Optogenetics, using a retrograde virus encoding channelrhodopsin injected into IC revealed a direct excitatory influence of neurones in auditory and motor cortices on firing in IC. Electrical stimulation of discrete cortical regions revealed that multiple non-auditory cortical regions have a direct monosynaptic excitatory influence on neurones in the IC which, in turn, activates local circuits increasing the firing probability of multiple neurones in the IC. This is the first evidence for circuitry by which auditory processing can be influenced at an early stage by activity in the sensory, motor and executive domains.

## Introduction

Increasing evidence suggests that identifying and responding to objects and events in the natural world involves interactions between the different senses, and the integration of sensory and motor processing in the brain (Kayser and Logothetis, 2007; Bizley and King, 2008; Brooks and Cullen, 2019; Bizley and Dai, 2020; Schneider, 2020; Lohse et al., 2022).

In perceptual processing, it is well established that behaviourally congruent information in one sensory domain can enhance perception in another, as illustrated by the benefit conferred by seeing the speaker’s lips in perceiving speech (Sumby and Pollack, 1954; Grant and Seitz, 2000; Peelle and Sommers, 2015), and the corollary that incongruent sensory information in one domain can disrupt perception in another as classically demonstrated by the McGurk effect (McGurk and Macdonald, 1976). For the most part, multi-sensory and sensory motor interactions are considered to occur at the level of the cerebral cortex. Numerous structural and functional studies in animals and humans have derived evidence for such interactions at primary and higher cortical levels, e.g., (Martuzzi et al., 2006; Bizley and King, 2008; Cappe et al., 2009; Jorge et al., 2018; Porada et al., 2019; Di Marco et al., 2021). Thus, widespread cortico-cortical projections likely mediate many cross-modal interactions. But in the case of the auditory system, the situation might be expected to be different. Unique among the sensory modalities, the auditory system features extensive pre-cortical (brainstem) processing networks, which potentially provide a substrate for multisensory integration much earlier in the pathway.

The inferior colliculus (IC, the auditory midbrain) represents the culmination of brainstem auditory processing and the source of almost all ascending auditory input to the thalamus and beyond (Ito and Malmierca, 2018; Rees, 2020). The IC receives afferent inputs from parallel pathways established in the brainstem as well as descending inputs from higher levels. That there are connections from the auditory cortex (AC) to the IC, particularly to its dorsal and lateral cortices, but including the central nucleus, is well established (Andersen et al., 1980; Coleman and Clerici, 1987; Feliciano and Potashner, 1995; Saldana et al., 1996; Bajo and Moore, 2005), but we have recently presented evidence that the IC also receives non-auditory cortical inputs. Using both anterograde and retrograde tracing, we demonstrated the existence of inputs to the IC from the visual (VC), somatosensory (SC), motor (MC) and prefrontal cortices (PFC) (Olthof et al., 2019). These inputs are glutamatergic, and they terminate on both GABAergic and non-GABAergic (putative glutamatergic) neurons in the IC. While these projections innervate all divisions of the IC, it is notable that they are also extensive in the central nucleus ICc (Olthof et al., 2019).

Consistent with our anatomical findings, several functional studies have suggested that the IC could be the recipient of inputs from cortical regions other than AC. Thus, modulation of VC has been demonstrated to influence the BOLD signal in the IC (Gao et al., 2015; Leong et al., 2018), while Groh and colleagues have shown that visual stimuli and saccade-related signals can evoke direct responses in the IC as well as modulate activity evoked by auditory stimuli (Groh et al., 2001; Porter et al., 2007). It has also been reported that self-generated movement modulates the responses of neurons in the IC and that these signals might be derived from efference copy from the motor cortex (Yang et al., 2020). Despite these indirect observations, to our knowledge there is no direct physiological evidence from neuronal recordings demonstrating the influence of inputs from non-auditory cortical regions on neuronal responses in the IC.

Here we report studies in which we have used both optogenetic and electrical stimulation in the rat to selectively activate areas of MC, SC, and PFC in combination with neuronal recording in the ICC. We present functional evidence for a direct influence of non-auditory cortical projections on neuronal activity in the ICC.

## Methods

### Animals

Male Sprague Dawley rats (Charles River) were housed in groups of 3 in two-storey individually ventilated cages under standard conditions (12h light/dark cycle lights on 7am; humidity 45-55 %, temperature 21-23 <C. The environment was enriched with cardboard tubes and wooden blocks/balls. Animals were allowed free access to food (rat chow SDS) and tap water. Animals were allowed to acclimatize for at least 3 days before experiments began.

### Virus injections

Rats were anaesthetised with isoflurane (5% in O_2_, 0.1 l/min) and ketamine/medetomidine (15/0.2 mg/kg, i.p.). The head was shaved, and the animal was moved to a sound-attenuated room and placed in a stereotaxic frame. The head was fixed using a custom-made tooth/palate bar and zygoma bars (WPI). The animal’s temperature was maintained with a homeothermic blanket set to 37 <C controlled by a rectal probe (Harvard). Anaesthesia was maintained with isoflurane in O_2_ delivered via a modified paediatric nasogastric cannula inserted into the nostril. The concentration of isoflurane was adjusted to maintain a surgical plane of anaesthesia. A small-animal pulse oximeter connected to a MouseVent G500 (Kent Scientific), clipped to one of the hind paws, was used to monitor arterial PO_2_ and heart rate.

The scalp was cut in the midline and the periosteum scraped to reveal the skull bones and sutures. Using a dental burr, a craniotomy was drilled (AP: -8.8 mm, ML: 2.5 mm from Bregma) to allow access to the right inferior colliculus. A multi-channel silicone electrode (10 mm single shank linear32 channel electrode, 100 µm pitch of recording sites, A1x65-10mm-100-177-A32 50, Neuronexus) was implanted into the IC. To avoid the superior and transverse sagittal sinuses, the electrode was implanted at an angle of 12 degrees to the vertical and 45 degrees from the sagittal plane (i.e., such that the electrode passed dorsoventrally from a rostral/lateral position to a caudal/medial position). The electrode was connected via a preamplifier (PZ2-32) to an RZ2 BioAmp (Tucker Davis Technologies, TDT) to amplify and record the neural signals. Voltages from 32 electrode channels were digitized at a sampling rate of 24414 Hz.

### Placement of IC recording electrode

To verify the location of the recording electrode in ICC we examined responses to sound stimuli. Pure tones at varying frequencies and amplitudes (75 ms duration, 512-16,384 Hz, 5-100 dB SPL) generated by an RZ6 Multi I/O Processor (Tucker Davis Technologies) were delivered to the animal’s ear canals via 10 cm lengths of Zygon tubing attached to MF-1 Multi-Field Magnetic Speakers (TDT). Frequency response area (FRA) plots of sound-evoked activity were constructed on-line (see Data analysis). Electrodes were considered to be in the ICC if they showed frequency-tuned, sound-driven, activity characterised by excitatory FRAs with a clear tonotopic progression (channels responding best to low frequencies located dorsally and channels responding best to high frequencies located more ventrally, see Figure 1). The electrode position was adjusted to optimise the number of channels located in the ICC and coordinates and depth recorded to guide the virus injection.

**Figure 1.**
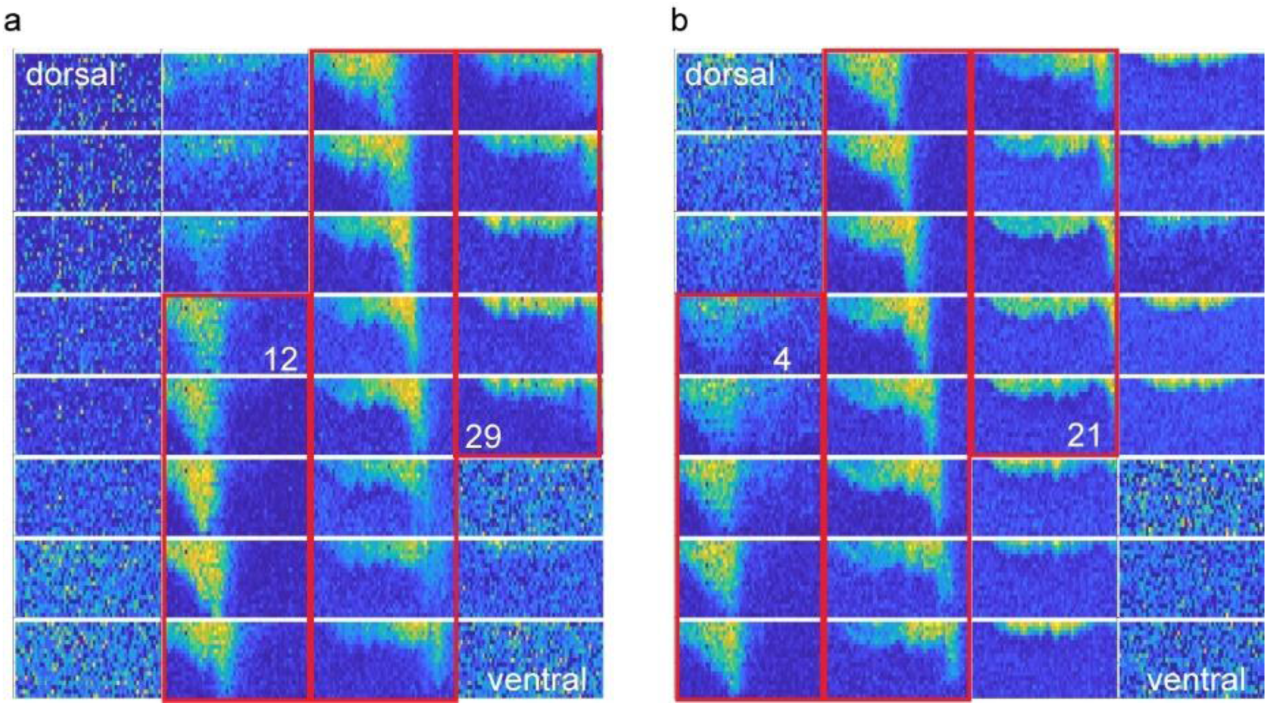
Example FRA plots from 32-channel electrodes implanted in the IC. Data from the 32 channels are arranged in 4 columns of 8 channels with upper left channel most dorsal (1) and lower right channel (32) the most ventral in the IC. Note that in example **a**, channels 12-29 (highlighted with red boxes) show a distinct tonotopic pattern of frequency response and would be considered to lie within the ICc. In example **b** channels 4-21 would be considered to lie within the ICc. Other channels lie outside the ICc

### Injection of virus

A glass capillary pulled to a fine tip was pre-loaded with a retrograde virus expressing both GFP and channelrhodopsin (ChR2) (pAAV-Syn-ChR2(H134R)-GFP (Addgene, 7 x 10^12^ vg/ml in PBS/NaCl/pluronic acid F-68) and fitted to an injection device (Nanoject, Drummond Scientific). The recording electrode was removed from the ICC and the glass pipette was implanted in its place. Virus was injected into the ICC (200-300 nl at three or four positions along the dorsoventral axis). To avoid leakage of virus out of the ICC, the pipette was left in place for 5 minutes after the final injection before being withdrawn. The animal was given postoperative analgesia (meloxicam, 1mg/kg, s.c.). The scalp was sutured (Vicryl 4.0) and the animal was removed from the stereotaxic frame and allowed to recover from the anaesthetic. Post-operatively, animals were housed in groups of 2 or 3 and were given soft diet for 1-2 days. They received an additional dose of meloxicam (1mg/kg, s.c.) on the day after surgery.

### Acute *in vivo* electrophysiological recording: optogenetic stimulation in virus injected rats

Acute electrophysiological recordings were conducted 12-40 days post virus injection. Animals were anaesthetized with urethane (0.5 g/kg i.p.) and, once the response to toe pinch was abolished, were injected with a mixture of fentanyl/midazolam (Hameln) (0.3/5mg.kg^-1^ i.m. in the *biceps femoris*). A tracheotomy was performed, and a polythene tube was inserted and tied into the trachea. Animals were moved to a sound-attenuated room and fixed in a stereotaxic frame (Kopf) on a homeothermic blanket, as above. To ensure good oxygen saturation throughout the experiment, the tracheal tube was connected to a pumped supply of O_2_ and the pumping rate was adjusted to be marginally quicker than the animal’s spontaneous breathing. Arterial PO_2_ and heart rate were monitored as above.

The scalp was cut in the midline and the periosteum scraped away to reveal the skull bones and sutures. The previous craniotomy over the IC was reopened. Further craniotomies were drilled to allow access to the auditory cortex and primary motor cortex (M1). For access to AC, the craniotomy was made medially and the stimulating electrode was implanted at an angle of approximately 20< mediolateral to the midline. The correct positioning of the electrode in AC was verified by the neural response to clicks.

To activate cortical neurones optogenetically, a glass fibre patch cable (M126L01 400 µm, 0.5 NA, Thorlabs, Ely, UK) was connected to a 470 nm LED (M470F3, Thorlabs, Ely, UK) with DC variable voltage LED driver (LEDD1B, Thorlabs, Ely, UK). The fibre was lowered to the surface of the brain. Ramped stimuli (0-5 V, over 5 or 10 s) were applied using a custom script in Spike2 (CED, UK). To examine local effects on cortical neurones, recordings were made using multichannel electrodes with four shanks and 8 channels per shank (Neuronexus, A4x8-5mm) implanted into the cortex close to the glass fibre. To examine the effects of optogenetic cortical activation on activity in the ICC, recordings were made using a linear 32-channel electrode (Neuronexus, A1x32-10mm) implanted into the IC. The location of the electrode within the ICC was optimised using the FRA measure as described above.

At the end of the acute electrophysiology experiment, animals were given an overdose of pentobarbitone (Euthatal, 400 mg.kg^-1^ i.p.) and transcardially perfused with heparinised 0.1 M phosphate buffered saline (PBS, ≍60 ml) followed by 4% paraformaldehyde (PFA, ≍60 ml) in PBS. The brain was removed and post-fixed in 4% PFA for 16-24 hours, cryoprotected in 30% sucrose in PBS, and rapidly frozen. Brains were sectioned on a freezing microtome and either immediately mounted or stored at -20 <C in antifreeze (30% ethylene glycol, 30% sucrose, 1% polyvinyl pyrrolidone (PVP)-40 in PBS) (Watson et al., 1986). Sections mounted on slides were counterstained with 4’, 6-diamidino-2-phenylindole (DAPI, Invitrogen) nuclear stain. GFP and DAPI were imaged utilising either a Nikon NiE widefield microscope equipped with an Andor Zyla 5.2 camera, a motorised stage and 10x/0.45NA air objective or a Zeiss Celldisoverer widefield microscope, equipped with a motorised stage and 20x/0.7NA objective.

### Acute *in vivo* electrophysiological recording: electrical stimulation in naïve rats

Naïve male Sprague Dawley rats (median 350g, range 240-386 g) were anaesthetized and prepared for surgery as described above. Craniotomies were drilled over the right IC and one or more of the following ipsi- and/or contralateral cortical regions: primary and secondary motor cortex (M1 and M2); somatosensory cortices-jaw region (S1Jaw), forelimb region (S1FL), hindlimb region (S1HL), barrel cortex, and medial prefrontal cortex (PFC), as delineated in the rat brain atlas of Paxinos and Watson (1998). Note: for access to barrel cortex, the craniotomy was made close to the midline and the stimulating electrode was implanted at an angle mediolateral to the midline.

A concentric bipolar stimulating electrode with poles separated by 500 µm (Clarke Electromedical) was implanted in the cortical region of interest targeting layer V. The two poles of the stimulating electrode were connected to a battery powered constant current stimulus isolator (Isoflex, AMPI, Israel). Electrical stimulation (triggered by the TDT system) were delivered as single cathodal square pulses of 100 µs duration at a rate of 2.5 Hz (i.e. one pulse every 400 ms) for a period of 40 s (100 sweeps, denoted as a stimulus ‘block’). To examine the threshold and current dependence of responses, the output of the stimulus isolator was varied between 0.3 and 8 mA for different blocks. Multiple cortical sites were stimulated in individual animals. Data were recorded from the 32 channels of the IC electrode during electrical stimulation of the cortex.

### Data analysis

To construct FRAs online, the signal was bandpass filtered (300-3000 Hz) and thresholded at ± 3 x the standard deviation of the average signal. Positive or negative voltage deflections greater than threshold were counted as spikes.

For all other analysis, we used a refinement of this spike thresholding script. The signal was filtered at 500-5000 Hz. A threshold was set at ± 2.5 x the standard deviation of the voltage signal across the 40 s data ‘block’ (excluding a period of 10 ms before and 80 ms after each stimulus to avoid the stimulus artefact and response). Where the voltage passed this positive or negative threshold, a spike was counted. To avoid double counting of spikes, where a positive and a negative deflection occurred within 1 ms, this was counted as a single, bipolar, spike. Peristimulus time histograms (PSTHs) were constructed by binning the spike data into 1 ms bins across the 400 ms sweep and summing the spikes in each bin over the 100 sweeps. The spontaneous firing rate was defined as the average rate in the last 40 ms of the sweeps. The total spikes presented in the PSTH is the sum of positive (only), negative (only), and bipolar spikes.

Having examined the nature of the excitatory and inhibitory responses (described below in Results), we designed custom scripts in Matlab to identify channels in the IC in which there was a response to electrical stimulation of the cortex, and to quantify the parameters of these responses. To avoid contamination of the response by the electrical stimulation artefact, responses were measured starting 3 ms from the time of electrical stimulation.

We used the following criteria to define excitatory and inhibitory responses to electrical stimulation: an excitatory response was defined as a group of at least 6, 1ms bins (consecutive or separated by 2 or fewer bins) in which the firing rate was at least 2 spikes above the spontaneous firing rate, and which contained at least 60 spikes in total. An inhibitory response was defined as at least 60 connected bins in which the moving average over 20 ms bins was at least 2 spikes below the spontaneous rate, and the overall average in that period was at least 4 spikes below the spontaneous rate.

Where group data are presented, the data from the 32 recorded channels were aligned on the basis of the low frequency end of the FRA (see Figure 1) so that, as far as possible, data from equivalent positions in the ICc (i.e. with similar frequency selectivity) are grouped together.

## Results

### GFP expressing retrograde virus injected in IC labels multiple cortical regions

Following injection of retrograde virus encoding GFP into the ICC, GFP was expressed in multiple cortical regions in the cerebral hemispheres both ipsilateral and contralateral to the injection. In coronal sections, retrogradely labelled cells were evident in deep layers of cortex. Cell somata were intensely labelled as were dendrites extending into more superficial layers. Fine axons could also be seen exiting the cortex and entering the white matter (Figure 2).

**Figure 2.**
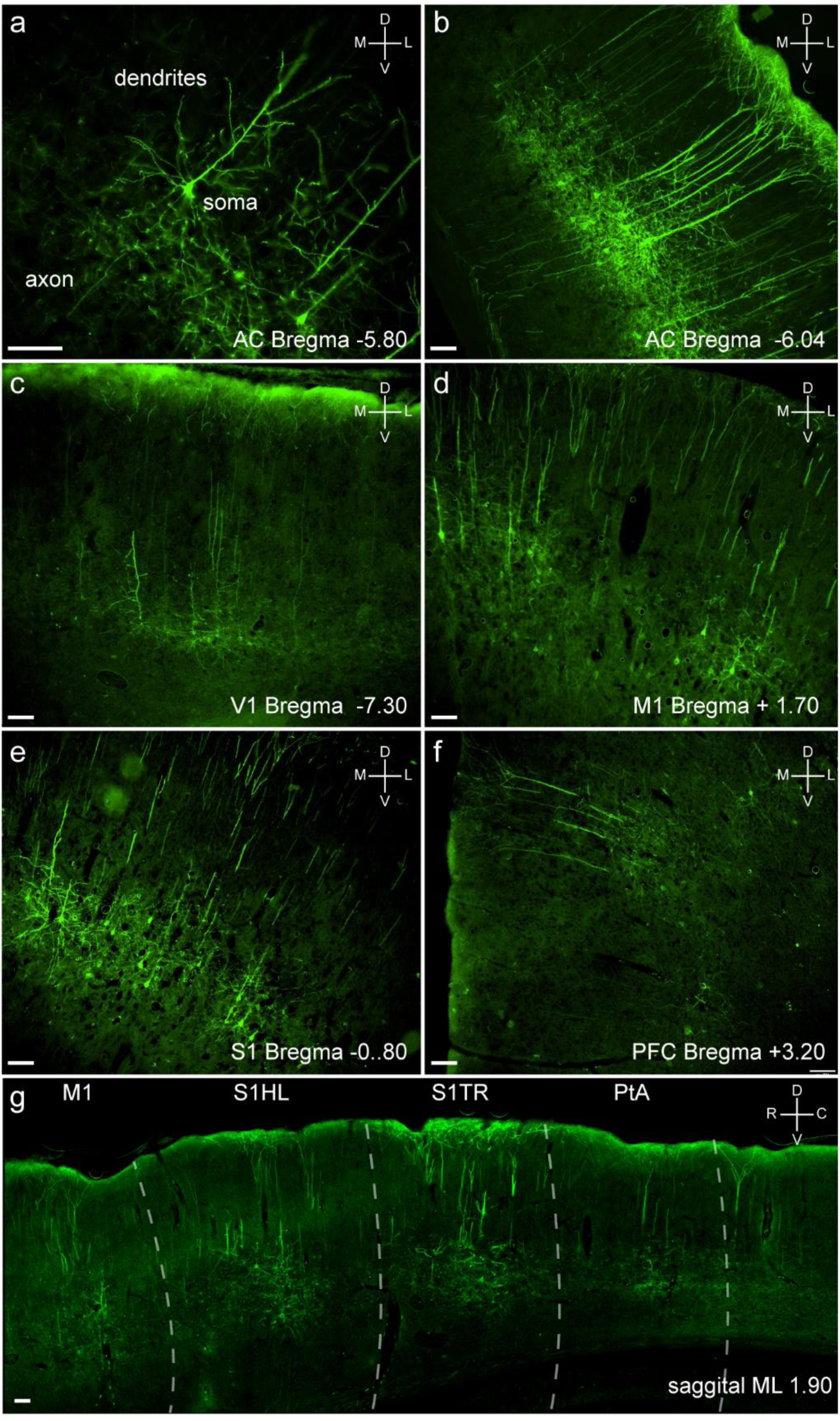
GFP labelling in regions of the cerebral cortex following injection of retrovirus in ICc. **a** pyramidal cell in AC showing GFP expression in soma, dendrites, and axon. **b** large numbers of GFP expressing pyramidal cells in AC layer V. **c** GFP expressing cells are also found in primary visual cortex, **d** M1, **e** somatosensory cortex and **f** PFC. **g** a sagittal section shows groups of GFP expressing cells in primary motor cortex, somatosensory hindlimb and trunk regions and parietal association cortex. Note that GFP-expressing cells are in small groups. Scale bar 100 µm.

Interestingly, although we injected virus at multiple points in the ICc to maximise coverage, GFP was expressed by relatively small groups of cells in restricted areas of cortex. The highest density of GFP expressing cells was in the ipsilateral AC, GFP expressing cells were also evident in the visual cortex (VC), somatosensory cortex (S1), motor cortex (M1) and prefrontal cortex (PFC) but at a lower density than in AC

### Channelrhodopsin expressing retrovirus injected into IC allows activation of cortical neurones

#### Local optogenetic activation of auditory cortex (AC)

The retrograde virus used also expressed channelrhodopsin (ChR2). To verify expression of ChR2 we examined the effect of blue light stimulation (470 nm) on neuronal activity in the cortex. To avoid desensitization, we stimulated the cortex with a ‘ramp’ of light of increasing intensity (Adesnik and Scanziani, 2010). Initially, we examined the AC as this was the region with the highest expression of GFP. In animals injected with the retrograde virus in the IC, application of blue light (ramped (0-5 V) stimuli of 5 or 10 s duration) to the AC evoked increases in neuronal activity which began a few hundred milliseconds after the light onset and ended as the light stimulus was switched off. Out of the eight virus injected animals tested, we were able to evoke activity in the auditory cortex in seven (see Figure 3a). In these animals, we were able to evoke responses on many electrode channels spanning a considerable area of AC. Responses were reproducible on repeated stimulation (Figure 3a) and the magnitude (number of channels activated or size/duration of response) was dependent on the light intensity (data not shown).

**Figure 3.**
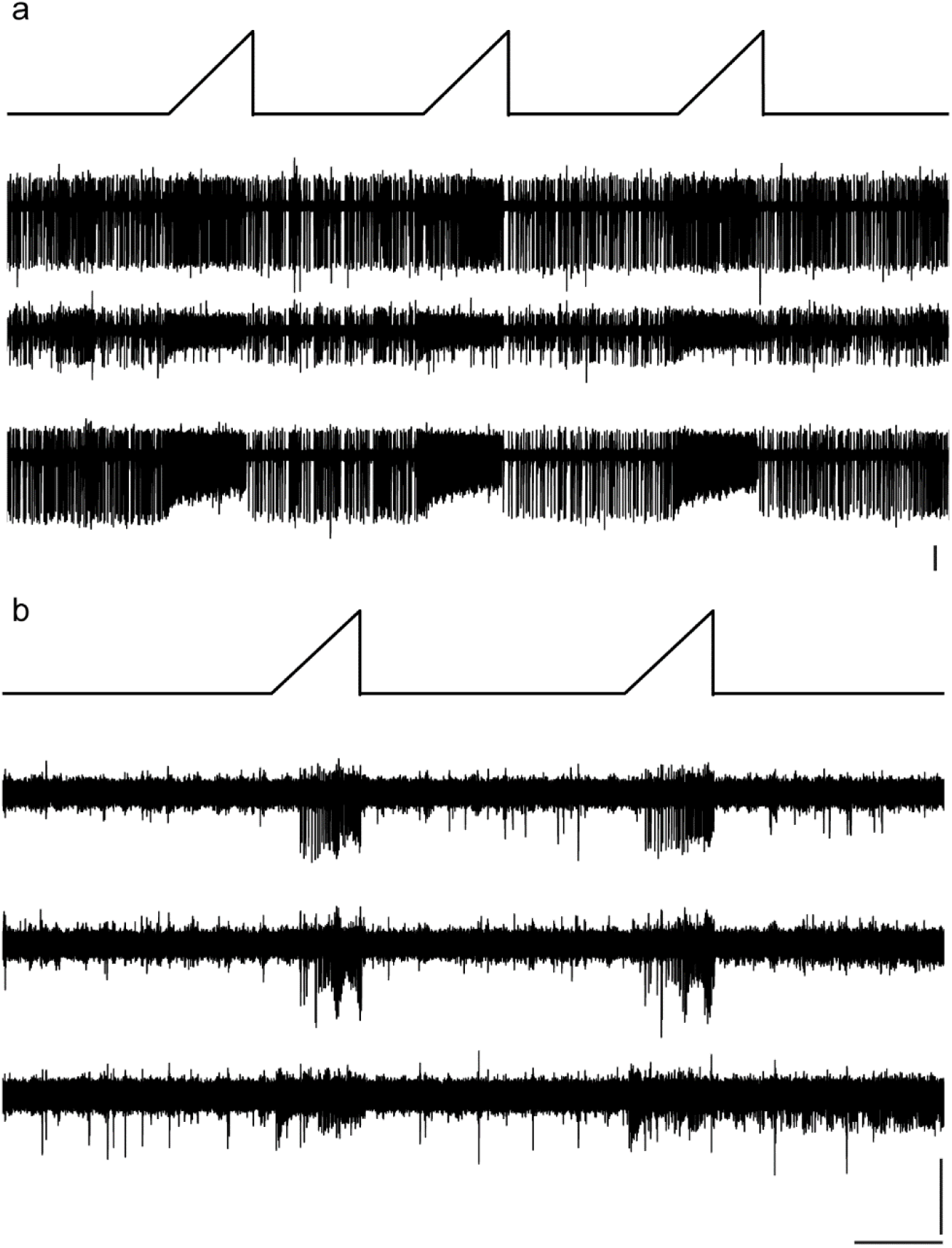
Optogenetic stimulation activates units in cortex. **a** shows the effect of three ramped light stimuli (0-5 V, 510 s) in AC on activity in three channels in ACx. **b** the effect of two ramped light stimuli (0-5 v, 5s) applied to M1 on activity on three channels in M1. Note that in each case the light stimulus increases the firing rate, that the responses to repeated stimuli are similar, and that multiple channels are affected. Top line in each part shows the ramped light stimulus (0-5V). Data were recorded on a 4 shank, 32 channel multielectrode probe. Vertical scale bars 50 mV; horizontal scale bars 10s.

#### Local optogenetic activation of M1/SS

Application of blue light to the M1 and somatosensory cortices also evoked an increase in activity in these areas (Figure 3b). Responses were like those evoked in the AC, indicating that the ChR2 was also functionally expressed in these regions. However, although we evoked activity in M1 and the border of M1/S1 in four animals, the regions where activation could be seen were small such that we failed to evoke activity at adjacent sites. Moreover, in several other animals, we failed to find regions in M1 or SS1 where the light stimulus evoked measurable increases in activity locally.

#### Optogenetic activation of AC and M1 evokes increased firing in ICc

Having verified that we could evoke firing activity in cortical neurones which project to the ICc (i.e., that contain ChR2 expressed by a retrogradely transported virus), we examined the impact of stimulating these neurones on activity in the ICC. As shown in Figure 4a, optogenetic stimulation of AC modulated firing activity in ICc

**Figure 4.**
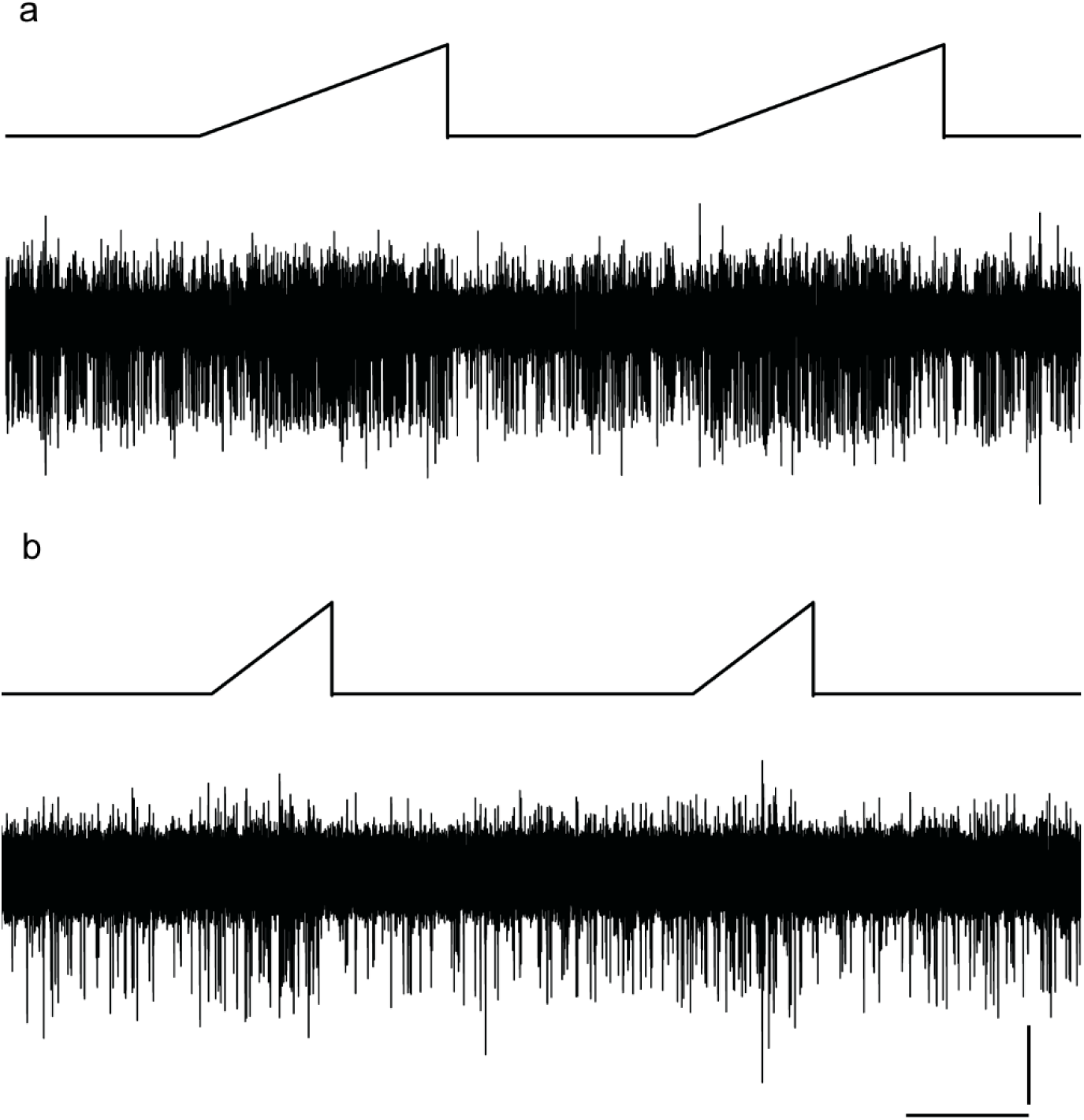
Optogenetic stimulation of cortex increases firing in ICc. **a** shows the effect of two ramped light stimulus (0-5 V, 10 s) applied to AC on activity on a single channel in IC_C_. **b** shows the effect of two ramped light stimuli (0-5 v, 5s) applied to M1 on activity on a single channel in IC_C_. Note that in each case the light stimulus increases the firing rate, that the responses to repeated stimuli are similar. Top line in each part shows the ramped light stimulus. Vertical scale bar represents 50 mV; horizontal scale bar is 10s.

The predominant response of neurones in the ICc to optogenetic stimulation of AC was an increase in the firing rate. The increase in activity was evident on several (usually adjacent) channels. However, this was observed in only 2 out of the 7 animals tested.

We also examined responses in ICc to optogenetic activation of M1. In one animal where we had seen local optogenetic activation of multiple channels in M1, we saw clear activation of several channels in ICc during optogenetic activation of M1 with the same parameters (Figure 4b). The increased firing rate occurred as the ramp of light increased and ceased when the light was off. Effects were seen on 6 adjacent channels in the ICc spanning 500 µm. However, we saw ICc responses to M1 activation in only one of 6 animals tested.

Given the highly localized, patchy, nature of the optogenetic labelling of cortex from the ICc, the relatively small numbers of neurones which could be activated using optogenetics, as well as the potential difficulty of locating the corresponding regions in the ICc where these fibres terminate, we chose to use electrical stimulation to further examine the impact of descending cortical connections on IC activity. Electrical stimulation also gave us the opportunity to assess the temporal relationship between firing in the cortex and in the IC.

### Electrical stimulation of primary motor cortex (M1) modulates activity in the inferior colliculus

We first examined the effects of electrical stimulation of primary motor cortex (M1) on activity in ICc. We chose this region because of its large size, ease of access, and the fact that in our tracing experiments, we had previously seen relatively large numbers of cells here retrogradely labelled from ICc (Olthof et al., 2020). Electrical stimulation of M1 (0.3-8 mA, 100 µs) evoked changes in the firing rate of units in the ipsilateral ICc. Effects were observed in 7 different animals (Table 1) and from multiple sites within M1.

**Table 1.** Stimulation sites evoking responses in ICc. In most cases multiple sites within a cortical region were tested. Not all sites were tested in all animals.

| Animal | M1 | M2 | Jaw | S1FL | S1HL | Barrel | PFC |
| --- | --- | --- | --- | --- | --- | --- | --- |
| ES03 | y |  |  | y |  |  |  |
| ES04 | y | y | y | y |  | y | y |
| ES05 | y | y | y | y | y | y | y |
| ES06 | y | y | y | y | y | y | y |
| ES07 | y | y | y | y |  | y | y |
| ES08 | y |  |  |  |  |  | y |
| ES09 | y | y |  |  |  | y | y |
| ES10 |  |  |  |  |  | y |  |

#### Excitation

Following electrical stimulation of M1, the most common response type in the ICc was an increase in firing activity. This was seen as both early fixed-latency events and longer latency more variable increases in firing described below.

##### Short fixed-latency event

Frequently, but not always, there was a short and fixed latency event around 4-5 ms after the peak of the stimulus artefact. In most cases, this early event could be discerned as a spike of relatively consistent shape and size which occurred following some or all of the individual stimuli (Figure 5a and 5b). For a fixed stimulus current (and fixed stimulation and recording locations), the latency of this early event was very consistent between sweeps (Figure 5a). The event was very evident when the responses to 100 stimuli were averaged (Figure 5c). Indeed, in some cases, this event was only discernible in the averaged signal. With increasing stimulus intensity, the probability of this spike event occurring in response to an individual stimulation increased meaning that the magnitude of the *averaged* signal deflection increased, although the size of the individual spike events did not change.

**Figure 5.**
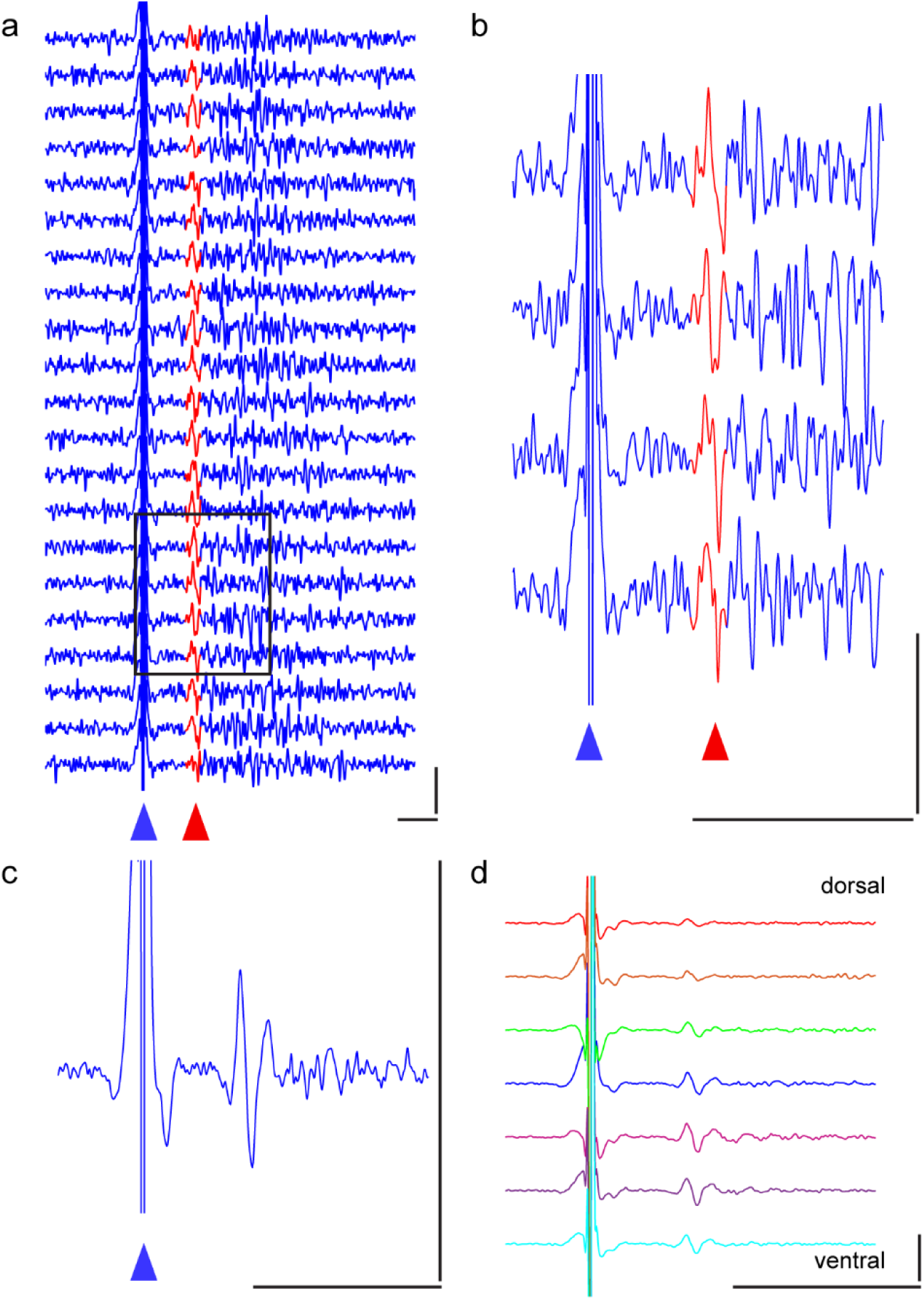
Electrical stimulation of M1 evokes a short latency firing event in ICc. **a** shows the response to 20 electrical stimulations (sweeps) recorded on a single channel in ICc. The stimulus consistently evokes a firing event with a short and fixed latency of around 5 ms (coloured red and indicated with a red arrow). The stimulus artefact is indicated with a blue arrow. **b** expanded view of four individual sweeps from the box in a. showing the relatively consistent shape and magnitude of the event. **c** shows the response averaged over 100 stimuli (40 s). **d** the short latency event is also evident on adjacent channels. Dark blue line channel shown in a, b, and c. Other coloured lines are data from channels located 300, 200 and 100 µm dorsal and ventral to this channel. Horizontal scale bars, 10ms; vertical scale bars 100 mV.

Often a short, fixed latency event was evident on several adjacent channels. The latency was very similar for responses on adjacent channels varying by less than 1 ms (Figure 5d).

##### Longer latency increase in firing

Irrespective of whether there was an early fixed latency event, electrical stimulation of M1 frequently evoked a relatively long period of increased firing activity in ICC. The latency and the duration of these later periods of increased activity varied but they usually began 5-10 ms after the stimulus artefact and continued for 10-30 ms (Fig 5a, 6ai and aii, bi and bii), but in some cases as long as 100 ms.

Although we could not definitively sort individual units from our multiunit activity recordings, it was clear from spike size alone, that multiple units contributed to this period of increased firing (Figure 6c). Examination of responses to individual stimulations showed that these longer latency firing events were more stochastic and that individual spikes occurred with a degree of temporal jitter. In some cases, we could tentatively identify a single unit with increased activity during this period (e.g. unit highlighted in Figure 6d) and could see that the latency of this individual unit to fire varied from sweep to sweep by several milliseconds.

**Figure 6.**
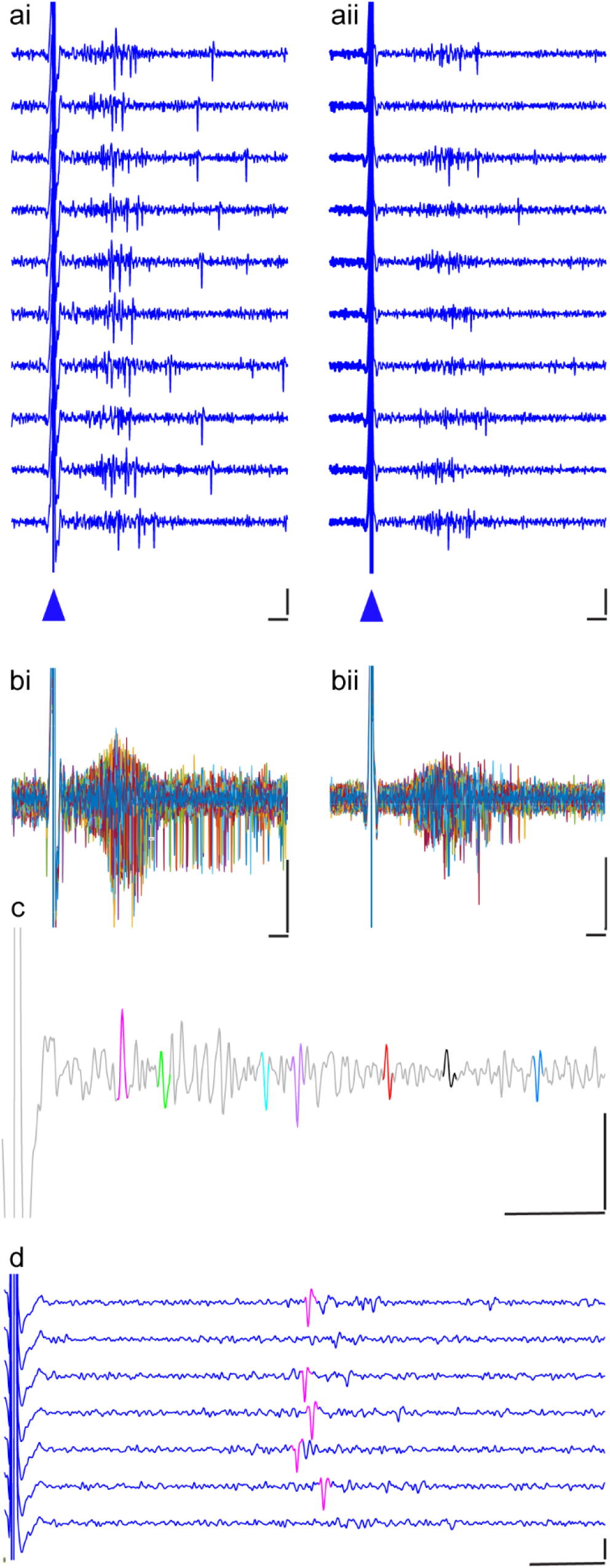
Example later excitatory responses in the ICc to electrical stimulation of M1. **ai** and **aii** show the responses to 20 stimulations recorded on a single channel in ICc in two different animals. Note the periods of increased activity following the stimulation artefact (indicated with a blue arrow). **bi** and **bii** show the same channels as ai and aii with all 100 sweeps overlaid and coloured differently. Note that the number of spikes varies between sweeps and that the response is longer in bi than in bii. **c** shows part of a single sweep recorded in ICc showing that the response to M1 stimulation involves multiple units, coloured differently. **d** shows that for a tentatively identified single unit (coloured magenta), there is temporal jitter in its occurrence and that some individual stimuli fail to evoke a spike in this unit. Vertical scale bars 100 mV, horizontal scale bars 5ms.

#### Responses are present on multiple channels-(excitation)

Although occasionally we saw responses to electrical stimulation on only one or two channels in the ICc, in most cases, the effect of electrical stimulation was evident on multiple adjacent channels within the ICc. Frequently we saw excitatory responses on 8 or 10 channels representing an overall linear distance in the ICC of 800-1000 µm (electrode pitch 100 µm) and occasionally on up to 24 channels. Although, because of the low resistance of the electrodes, in some cases the same individual units were recorded on two adjacent channels, comparison of the spike timings in the signal recorded from adjacent channels revealed that many different units were activated by the stimulus (Figure 7). Moreover, it is highly unlikely that the same units would be recorded over distances greater than 200 µm, suggesting that a large number of unts in the ICc were activated.

**Figure 7.**
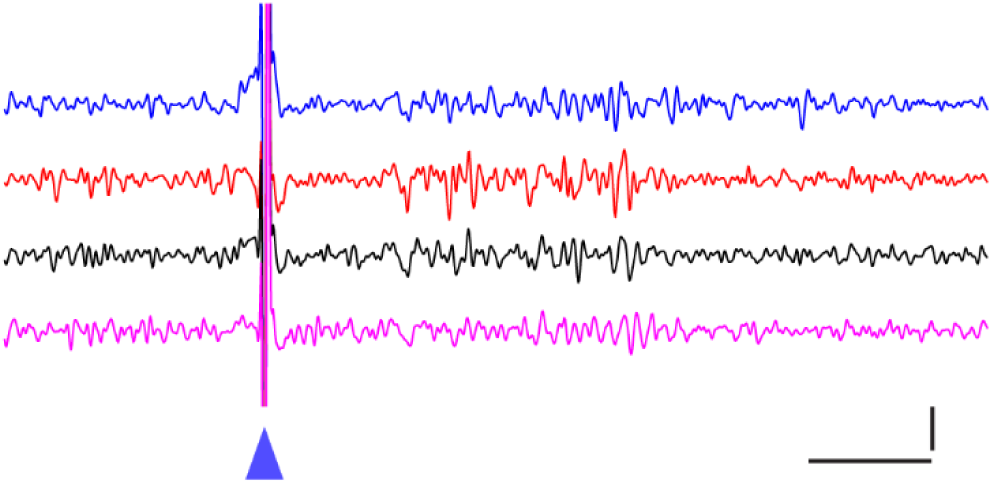
Activity on four adjacent channels in the ICc evoked by a single electrical stimulation of M1. Blue (dorsal), red, black and magenta (ventral) traces are from adjacent channels in the ICc spaced 100 µm apart. Note that there is increased activity on all four channels between 5 and 25 ms after the stimulus (blue arrow), that different shapes and sizes of spike can be seen on individual channels, and that the some of the spikes differ in timing between channels. Vertical scale bar 100 mV, horizontal scale bar 5 ms.

#### Inhibition

In some animals at some stimulation/recording sites, electrical stimulation of M1 evoked inhibition of firing in the ICc (Figure 8). In general, inhibition occurred with both a longer latency, and a longer duration than the previously discussed excitatory responses. This type of inhibition occurred independent of whether excitation was observed on that particular channel. Very occasionally, we observed inhibition with a short latency and duration (e.g., Figure 8c). Note that due to the low spontaneous firing rate in the absence of auditory stimulation, the inhibition was always of limited magnitude.

**Figure 8.**
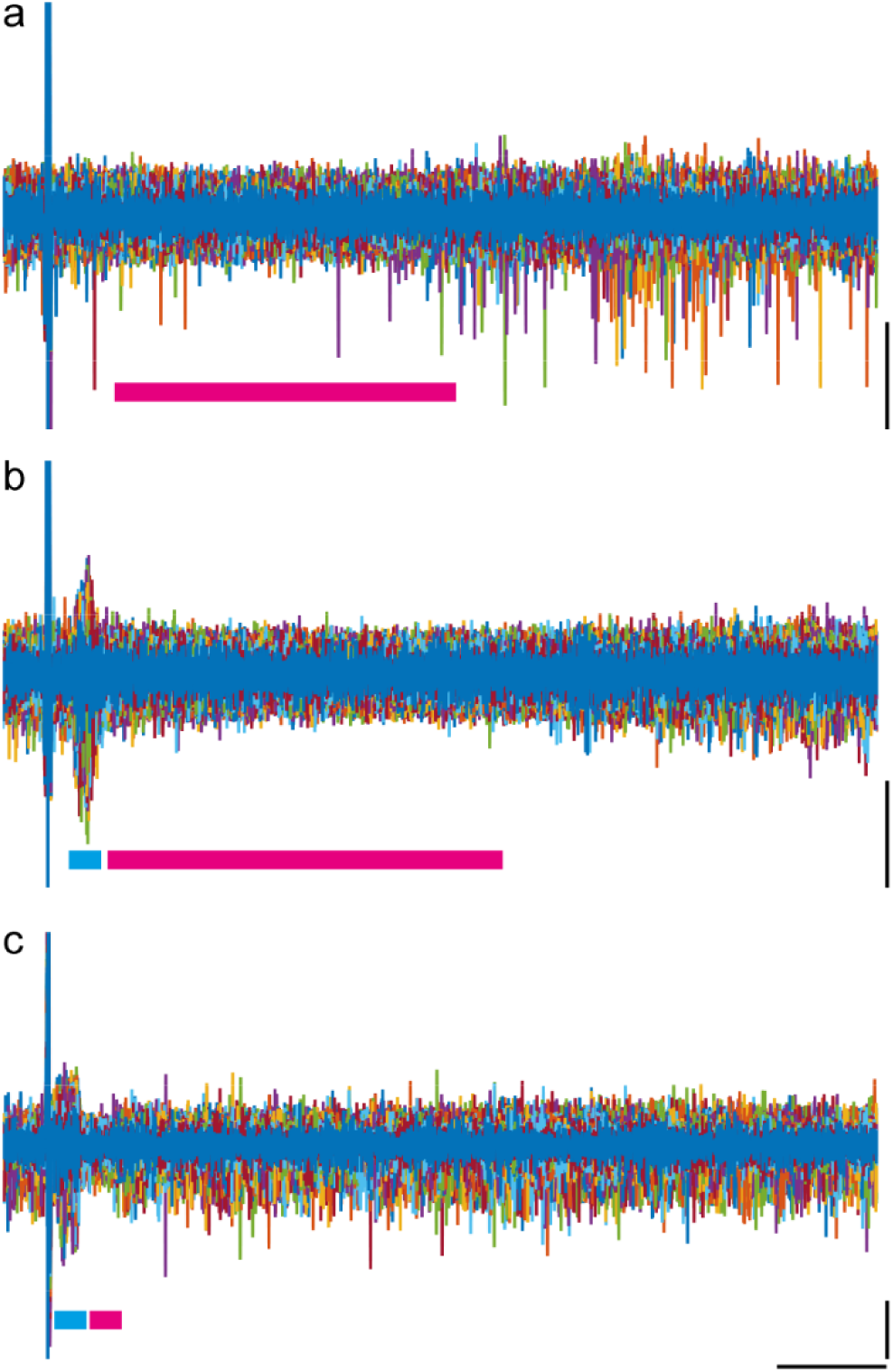
Activity on individual channels in the ICc during electrical stimulation of M1. Data are from 100 individual sweeps overlaid. Each sweep is coloured differently. The large deflection on the left is the electrical stimulation artefact. Coloured bars represent periods of excitation (cyan bar) and inhibition (magenta bar). **a** shows an example where, in the absence of any short-latency excitatory response, there is a long duration period of inhibition. **b** an example where there is a substantial increase in firing activity with short latency and relatively low jitter, which is followed by a later period of inhibition. **c** an example in which a short period of inhibition follows a short latency excitation. Note that this channel exhibited an unusually high spontaneous firing rate. Magenta bars indicate periods of inhibition, blue bars indicate periods of excitation. Vertical scale bars 50 mV, horizontal scale bar 50 ms.

In summary, electrical stimulation of M1 evoked changes in firing rate in ICc. Most responses were excitatory. In some cases, there was a clear short-latency event with highly consistent timing. More often a later period of increased firing activity of duration from approximately 10 to 50 ms, involving multiple units with marked temporal jitter. Inhibition was seen on a few occasions. We confirmed the biological nature of these responses by verifying that when we applied electrical stimulation post mortem, only the stimulus artefact was apparent.

### Responses occur from multiple sites in M1

In the rat, M1 is extensive in both mediolateral and rostrocaudal dimensions. To determine whether the connections between M1 and ICc are spatially restricted (as suggested by the viral labelling), we stimulated at multiple sites within M1, in most cases keeping the recording electrode in the same position in the ICc. While we were able to elicit responses in the ICc from multiple sites within M1, we also noted that there were many locations in M1 from which we did not elicit measurable responses at the recoding site in ICc. Thus, responses were highly localized, such that electrode sites within 200 µm of an ‘active’ stimulation site, sometimes failed to elicit any response at a particular recording site in ICc (Figure 9).

**Figure 9.**
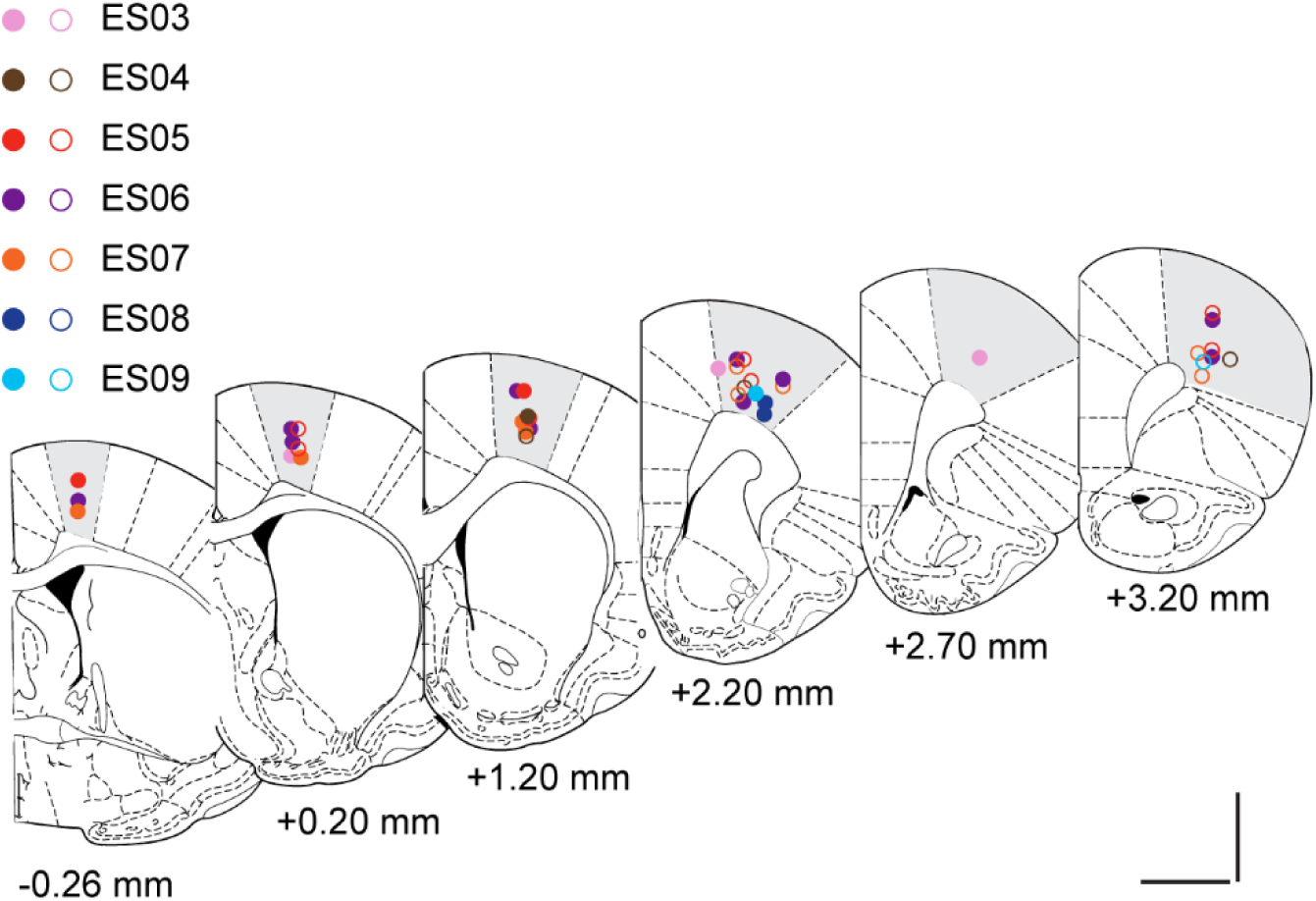
Stimulation sites in M1. Grey shading indicates areas denoted as M1 in the Rat Brain Atlas (Paxinos and Watson, 1988). Filled symbols indicate sites from which measurable (excitatory) responses were evoked in ICc. Open symbols show stimulation sites from which we failed to detect a response at currents up to 8 mA. The different colours denote different animals. Distances are anterior/posterior to Bregma. Scale bars 2mm.

To examine the characteristics of the response of ICc neurones to stimulation of M1, we pooled data from all M1 stimulation site/ICc recording sites in all animals in which we had observed measurable (early and/or later) responses. Animals and stimulation/recording sites from which current up to 8 mA failed to elicit a response are excluded. The resulting data set consisted of 22 stimulation site/recording site pairings in 7 animals where we successfully matched stimulation and recording sites that were interconnected.

First, we examined the early fixed latency event. This was present (on at least one channel) in 3/7 animals at 15 stimulation/recording site pairings. Interestingly, as the stimulus intensity was increased, the latency of the early event decreased (Figure 10a and b) and the number of channels showing an early event increased (Figure10b). The magnitude of the averaged response also increased (Figure 10a), but notably, however, on individual sweeps it was an all or none event of invariant size. Thus, the difference in magnitude in the averaged response represents an increase or decrease in the number of events. In cases where there was an early event on multiple channels, we found that (for a fixed current) the latency varied very little from channel to channel.

**Figure 10.**
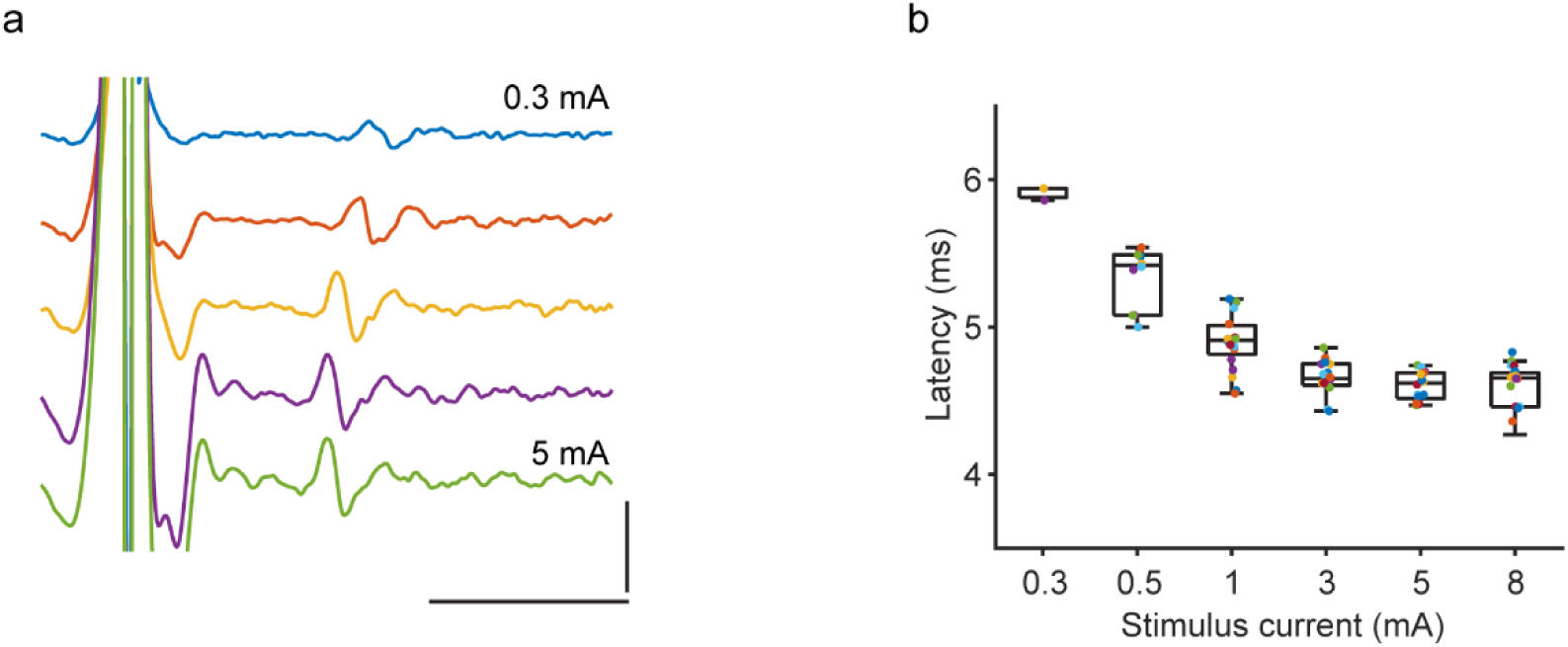
Increasing stimulus intensity in M1 decreases the latency of the early fixed latency event in ICc. **a** an example showing that increasing current from 0.3 mA (blue) to 5 mA (green) results in a progressive decrease in the latency. Note that the size of the averaged event increases with increasing current reflecting the fact that at higher currents, the event occurs with greater probability. Vertical scale 50 mV, horizontal scale 5 ms. **b** shows group data from 15 M1 stimulation sites in 3 animals in which between 2 and 5 different stimulus currents (0.3-8 mA) were tested. Note that there is a decrease in latency with increasing stimulus current. Each spot represents a single channel in ICc showing a short fixed-latency event. Different colours represent different stimulation/recording site pairs.

Next, we examined the late excitatory response evoked by electrical stimulation of M1. The latency of this response was generally between 5 and 10 ms from the stimulus and the latency showed no noticeable stimulus-current dependence and did not vary with recording channel for those channels adjudged to be within the ICc. However, the magnitude of this response (sum of the evoked spikes) was current dependent. Most responses emerged following stimulation at 1 mA and were near maximal at 3 mA, such that the current response curve was very steep. However, in a few cases, responses were seen at 0.3 or 0.5 mA and increased gradually with increasing current (up to 8 mA). It was notable that the number of channels on which a later excitatory response could be seen increased with increasing current. However, for any given current, the magnitude of the response diminished gradually for recording locations both dorsal and ventral to the most affected channel (Figure 11).

**Figure 11.**
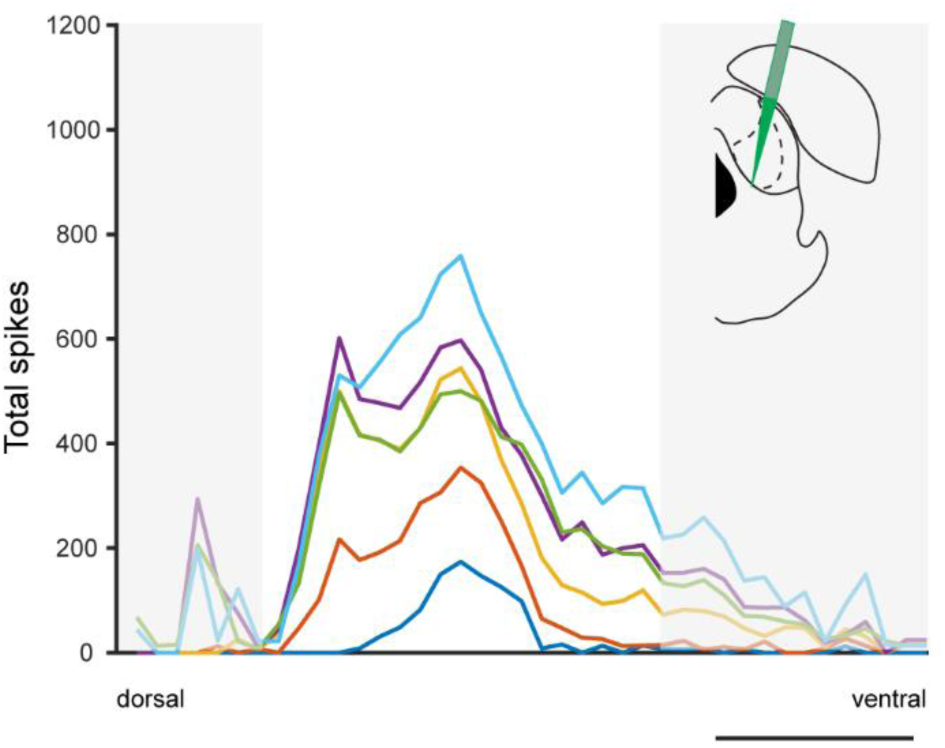
Increasing stimulus intensity increases the magnitude of the response and the number of channels affected by stimulation of M1. Stimulation currents: blue 0.3 mA, orange 0.5 mA, yellow 1 mA, purple 3 mA, green 5 mA, blue 8 mA. Data are from between 3 and 30 stimulation sites in 7 animals FRAs aligned: data greyed out are from channels dorsal (left) and ventral (right) to the ICc. Inset shows approximate position of the recording electrode in the IC. Scale bar 1 mm

### Electrical stimulation of secondary motor cortex (M2) also evokes excitatory responses in IC

Following electrical stimulation of M2, we also saw excitatory responses in 5 animals at 14 stimulation/recording site pairings (Figure 12). Early fixed latency events were seen in 2 animals (7 stimulation/recording site pairings). These events were similar in latency and showed similar current-dependence to those evoked by stimulation of M1 (Figure 13a), although a decrease in latency from dorsal to more ventral ICc was noted (data not shown). Irrespective of whether an early event was seen, electrical stimulation of M2 commonly evoked a period of increased activity beginning at around 5 ms (5 animals, 14 stimulation/recording site pairings). Responses occurred on multiple channels in ICc with more dorsal channels affected at the lowest currents and to the greatest degree (Figure 14a). With increasing current, the size of the response on individual channels was increased and responses were seen on more channels-in particular channels located more ventrally (Figure 14a). The later excitatory responses evoked by electrical stimulation of M2 were of similar latency and magnitude to those evoked by stimulation of M1 (c.f., Figures 10 and 13a, 11 and 14a). Occasionally there was a period of inhibition (data not shown).

**Figure 12.**
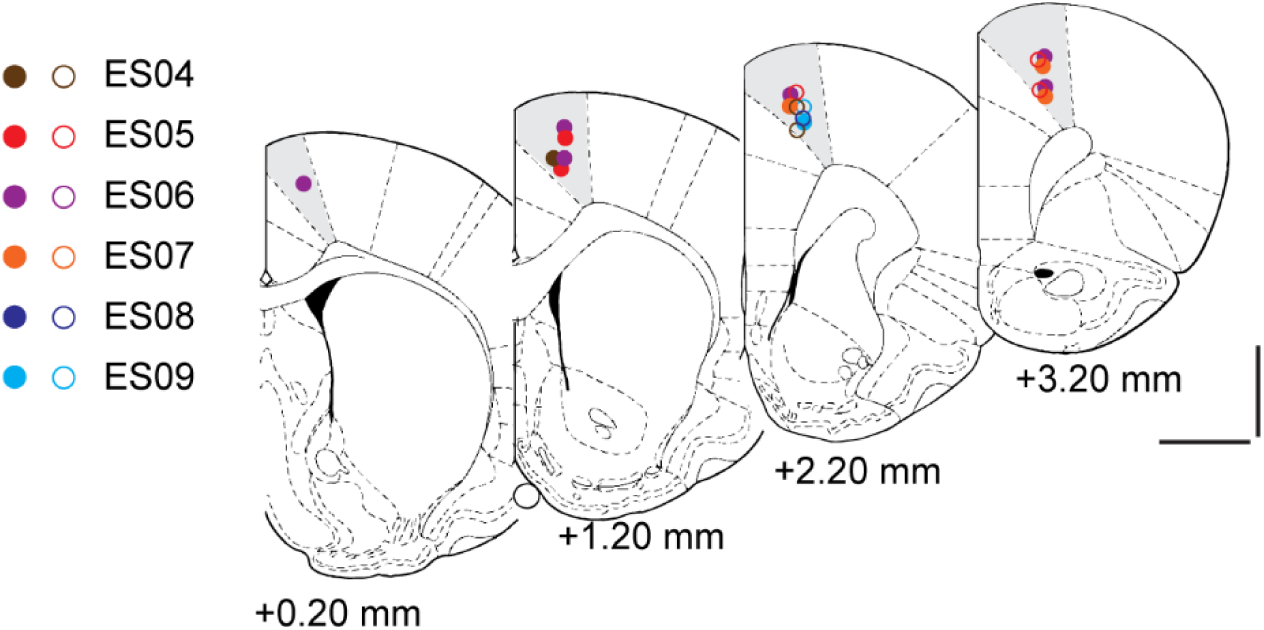
Electrical stimulation sites in M2. Data are from 6 animals. In 5 animals, electrical stimulation of at least one site in M2 evoked an excitatory response in ICc. In one animal (ES08) only one site in M2 was tested and stimulation at this location failed to evoke a response in ICc. Filled symbols represent sites from which an excitatory response was evoked. Open symbols represent sites from which no response was evoked. Different colours represent the different animals. Distance are anterior to Bregma. Scale bars 2mm.

**Figure 13.**
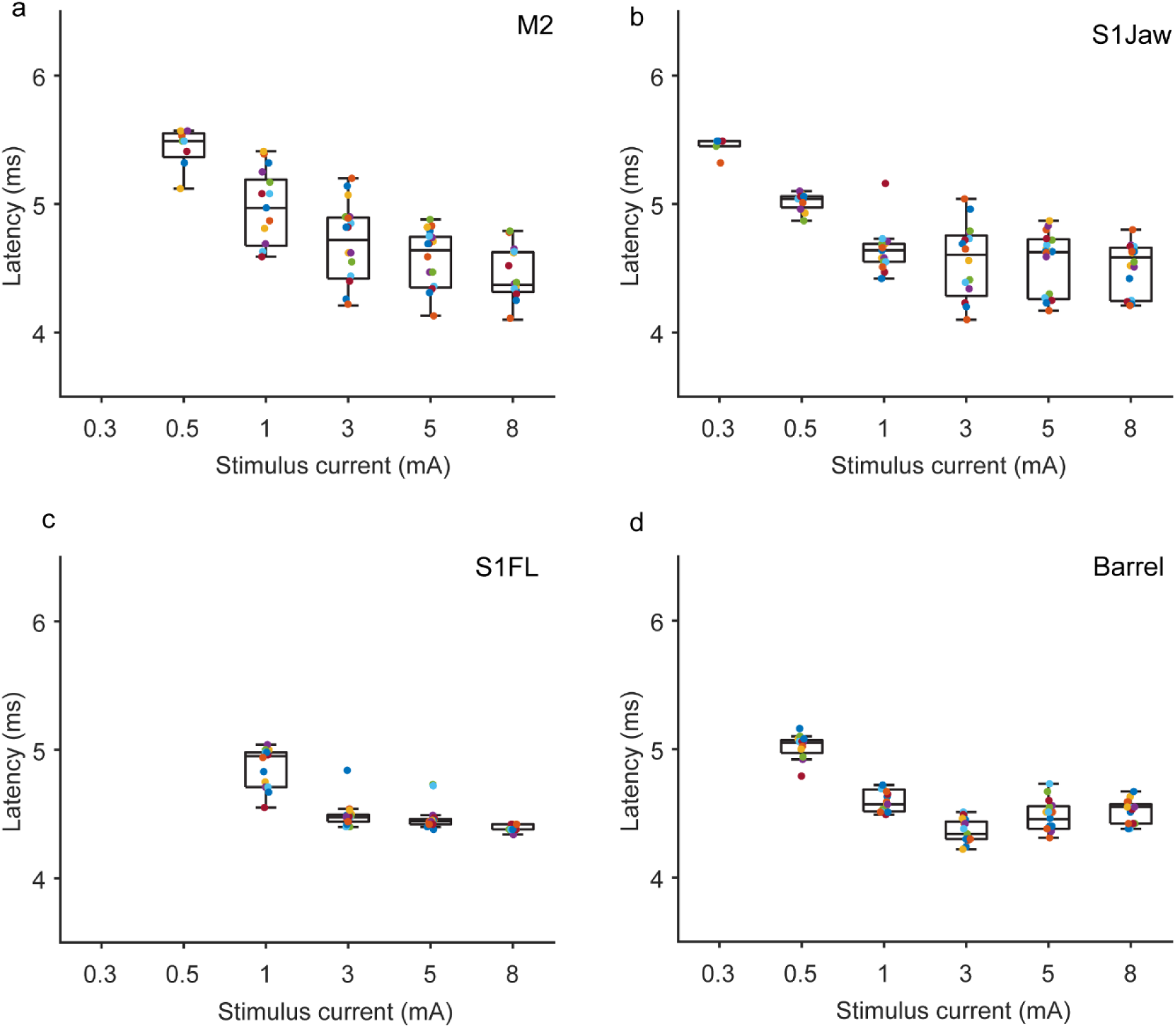
Electrical stimulation of **a** M2, **b** SIFL, **c** S1Jaw, and **d** barrel cortex evokes a short, fixed latency event in ICc. Data are from multiple stimulation sites in 2 animals (M2, S1Jaw, S1FL), or 3 animals (barrel cortex) in which between 2 and 5 different stimulus currents were tested. Different coloured spots represent different stimulation/recording site pairs. Note that there is a general decrease in latency with increasing stimulus current.

**Figure 14.**
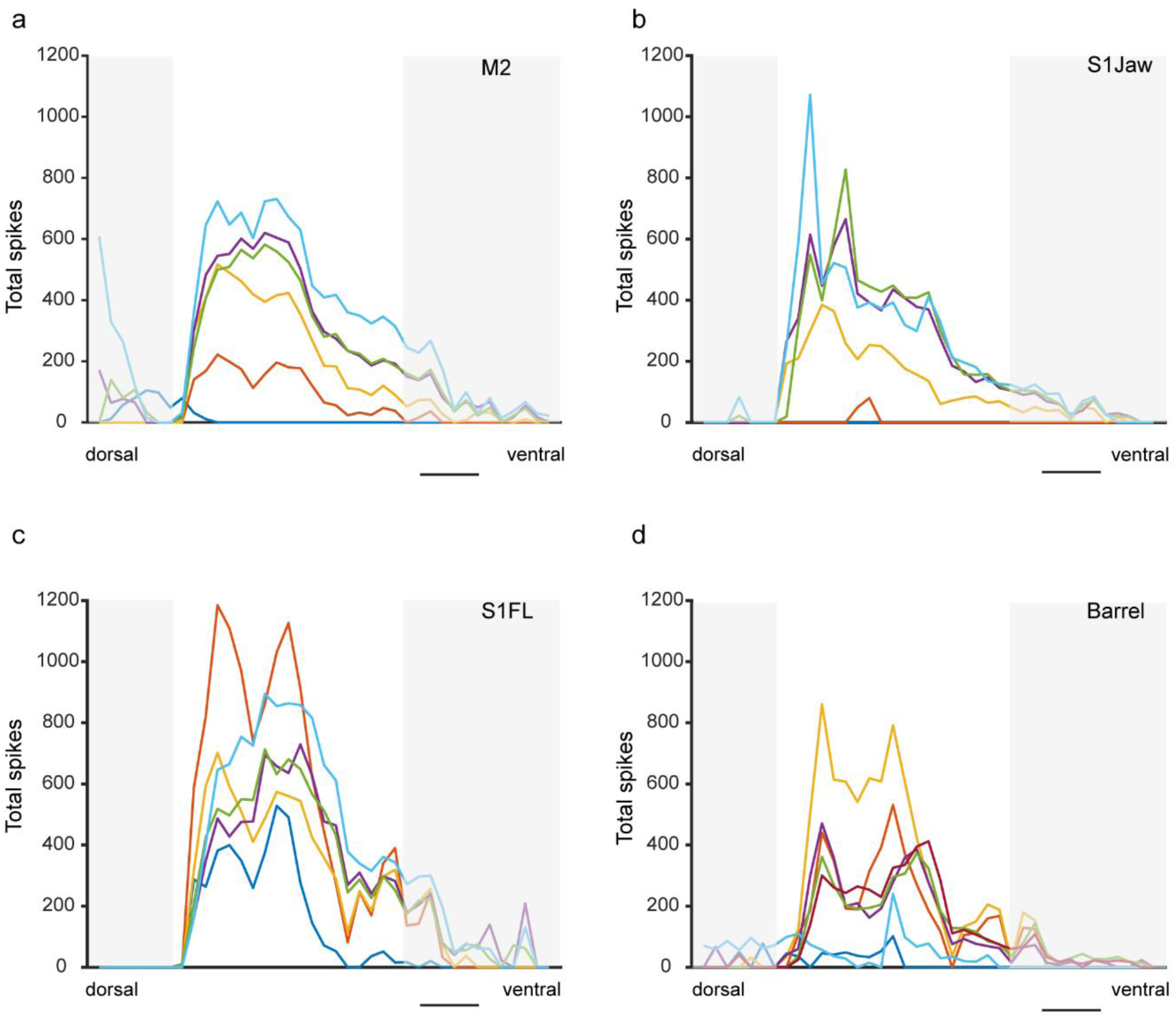
Increasing stimulus increases the magnitude and the number of channels on which a later excitatory response is seen following stimulation of **a** M2, **b** S1Jaw, **c** S1FL, and **d** barrel field. Stimulation currents: blue 0.3 mA, orange 0.5 mA, yellow 1 mA, purple 3 mA, green 5 mA, blue 8 mA. Data are summed PSTH in the period defined as a ‘response’-see Methods and are from multiple stimulation sites in 5 animals (M2, S1FL), 4 animals (S1Jaw) or 6 animals (barrel) in which between 2 and 5 different stimuli were tested. Note that with increasing stimulus current there is a general increase in the response magnitude (responses evoked by stimulation at 0.5 mA in S1FL and 1 mA in barrel field do not follow this trend). FRAs aligned: data greyed out are from channels dorsal (left) and ventral (right) to the ICc. Scale bar 500 µm.

### Electrical stimulation of somatosensory regions evokes increased firing in ICC

In addition to the motor areas M1 and M2, we examined the effect of electrical stimulation of several ipsilateral somatosensory cortical regions (S1Jaw, S1Forelimb (S1FL), S1Hindlimb (S1HL), barrel field) with several different cortical sites tested in each animal (see Table 1). As shown in Table 1, we recorded excitatory responses in ICc in response to stimulation of somatosensory cortical sites in multiple animals: S1Jaw (n=5), S1FL (n=6), S1 HL (n=2), and barrel field (n=6).

#### Somatosensory jaw (S1Jaw)

We observed excitatory responses to stimulation of the somatosensory jaw area (S1Jaw) in 4 animals (13 stimulation/recording site pairs). In one animal with the recording electrode in a single position in IC, we failed to elicit a response from 3 different stimulation sites in S1Jaw (Figure 15), suggesting that connections between cortical areas and the IC are spatially restricted.

**Figure 15.**
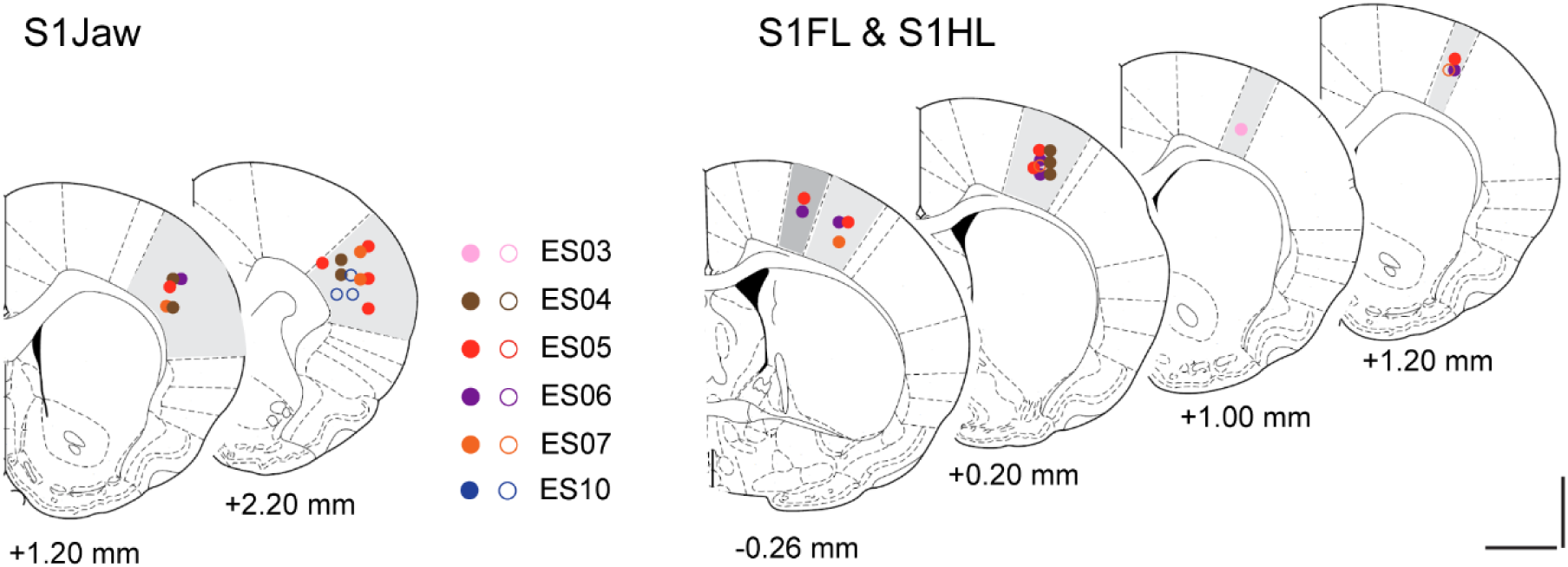
Electrical stimulation sites in S1FL, S1HL and S1Jaw regions. Grey shading indicates areas denoted as S1Jaw, S1FL, and S1HL (darker grey) (Paxinos and Watson, 1986). Filled symbols indicate sites from which measurable (excitatory) responses were evoked in ICc (0.5-8 mA). Open symbols denote sites from which there was no measurable response in ICc at currents up to 8 mA. Different colours represent different animals. Distances are anterior/posterior relative to Bregma. Scale bars 2mm.

Following stimulation of the S1Jaw region, early fixed latency events were seen in 2 animals (4 stimulation/recording site pairs). The latency of this event decreased with increasing current (1-8 mA) (Figure 13b) but there was no systematic variation in latency at sites across the ICc.

As in the motor regions, irrespective of whether we saw an early event, stimulation of S1Jaw evoked a later excitatory response which took the form of an increase in firing probability of multiple units with temporal jitter. This type of response was observed at all 13 stimulation/recording site pairs tested in 5 animals. The latency of this late response was relatively constant across recording sites and did not vary systematically with current (data not shown). However, the magnitude of the response and the number of channels affected did increase with increasing stimulation current (Figure 14b). It was notable that both the early and later responses to stimulation of the S1Jaw area appeared strongest in the most dorsal parts of the ICc (Figure 14b).

#### Somatosensory forelimb (S1FL) and hindlimb (S1HL)

Electrical stimulation of S1FL also evoked excitatory responses in the ICc (5 animals at 12 stimulation/recording site pairings). Responses could be triggered from multiple sites within S1FL, although we also found regions from which stimulation did not evoke a response (Figure 15). In 3 animals (6 stimulation/recording site pairings), electrical stimulation evoked a short, fixed latency event. The latency decreased and the number of channels on which a response was seen increased with increasing current (Figure 13c).

In 5 animals at 12 stimulation/recording site pairings we saw a late increase in firing activity beginning 5-10 ms after the stimulus which involved multiple units with temporal jitter. The latency was not current-dependent and there was no systematic variation in latency with recording site location within the ICc (data not shown). The magnitude of the later response and the number of channels affected was generally dependent on the stimulus intensity (Figure 14c).

Electrical stimulation of S1HL was tested in 2 animals as shown in Figure 15. In both cases an excitatory response was evoked in ICc (data not shown).

#### Barrel cortex

Responses in the ICc to stimulation of the barrel field were observed in 6 animals at 11 stimulus/recording positions. Responses to stimulation of the barrel cortex were like those evoked by stimulation of other somatosensory areas. In some cases, there was an early spike event of fixed latency (3 animals at 5 stimulation/recording pairings): the latency generally decreased with increasing current (Figure 13d) but did not vary with recording position).

We also saw a later excitatory response to electrical stimulation of barrel cortex. This began with a latency of 5-10 ms which was independent of current and location within the ICc. The magnitude of the response generally increased with increasing current (Figure 14d).

### Electrical stimulation of Prefrontal Cortex evokes increased firing in IC

Responses in the ICc were also observed following electrical stimulation of the PFC (11 anterior posterior stimulation sites in 6 animals). Because of the dorsoventral orientation of this region, we were able to stimulate all subregions sequentially beginning with the most dorsal (cingulate Cg) by lowering the stimulating electrode progressively through the prelimbic (PrL), infralimbic (IL) and dorsal peduncular (DP) areas. Electrical stimulation evoked excitatory responses comprising a period of increased firing activity which sometimes preceded by a short fixed-latency spike. On a very few occasions we saw a short latency event in the averaged signal over 100 sweeps without any longer latency increase in firing.

As in other regions, the response magnitude and number of channels affected was dependent on the stimulus current applied, albeit with a very steep stimulus-response curve. It was notable that, in most cases, channels which showed activation from Cg also showed activation from other PFC subregions. This was the case for both short latency (e.g. inset Figure 16) and longer latency components (Figure 17). There were no obvious differences in the response profile evoked by stimulation of the different PFC subregions and, for a given current, the magnitude of response to stimulation IL and DP subregions was marginally greater than the more dorsal Cg and PrL subregions (Figure 17).

**Figure 16.**
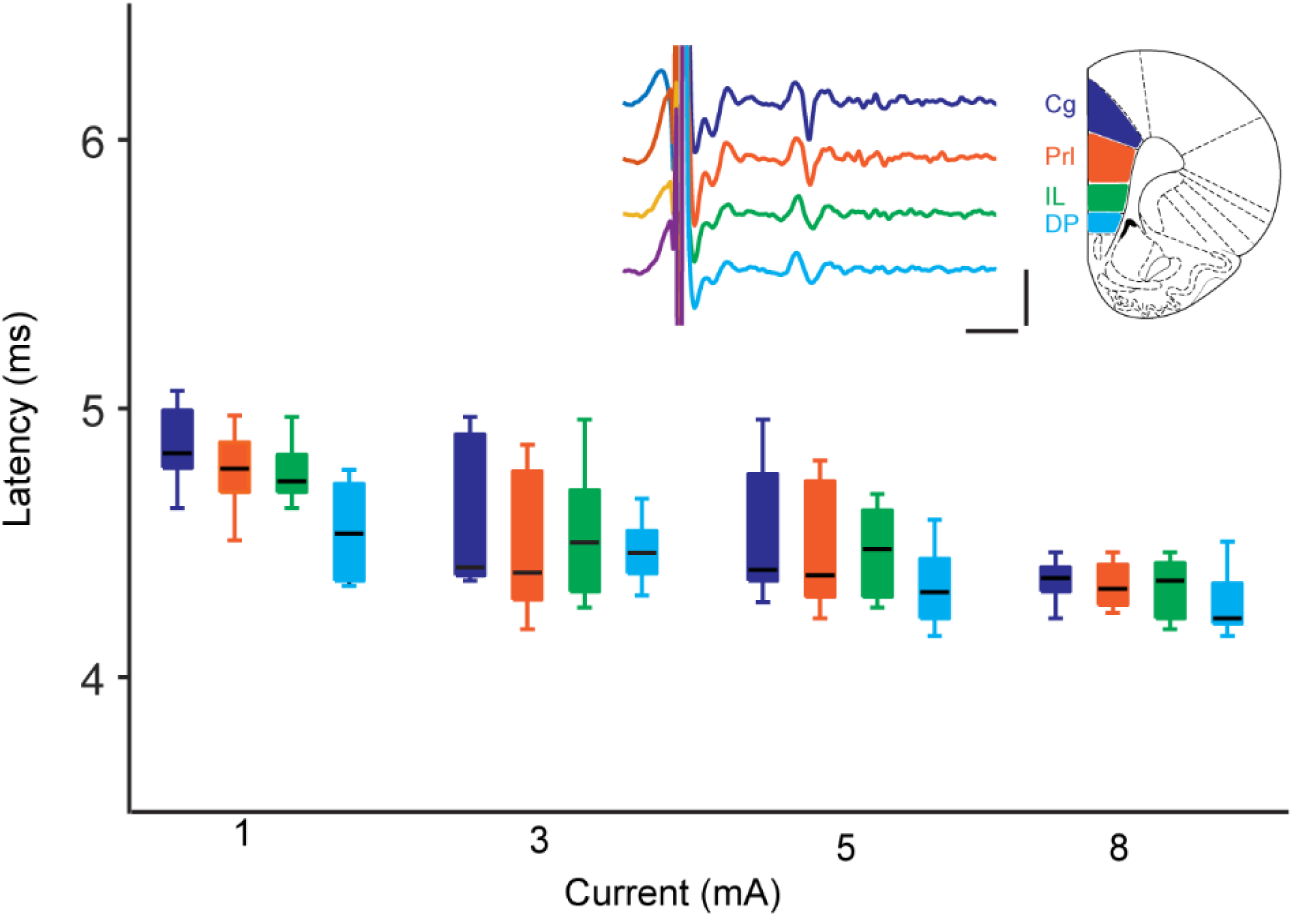
The latency of the early fixed latency event in IC_C_ evoked by stimulation of PFC subregions. Cg (dark blue), PrL (orange), IL (green), DP (light blue). Data are from multiple stimulation sites in 4-6 animals. Note that there is no consistent difference in latency between subregions but that the latency decreases with increasing current. Line: median; box: IQ range, whiskers: extreme data points. Inset shows example data. Vertical scale bar 100mV, horizontal scale bar 2 ms.

**Figure 17.**
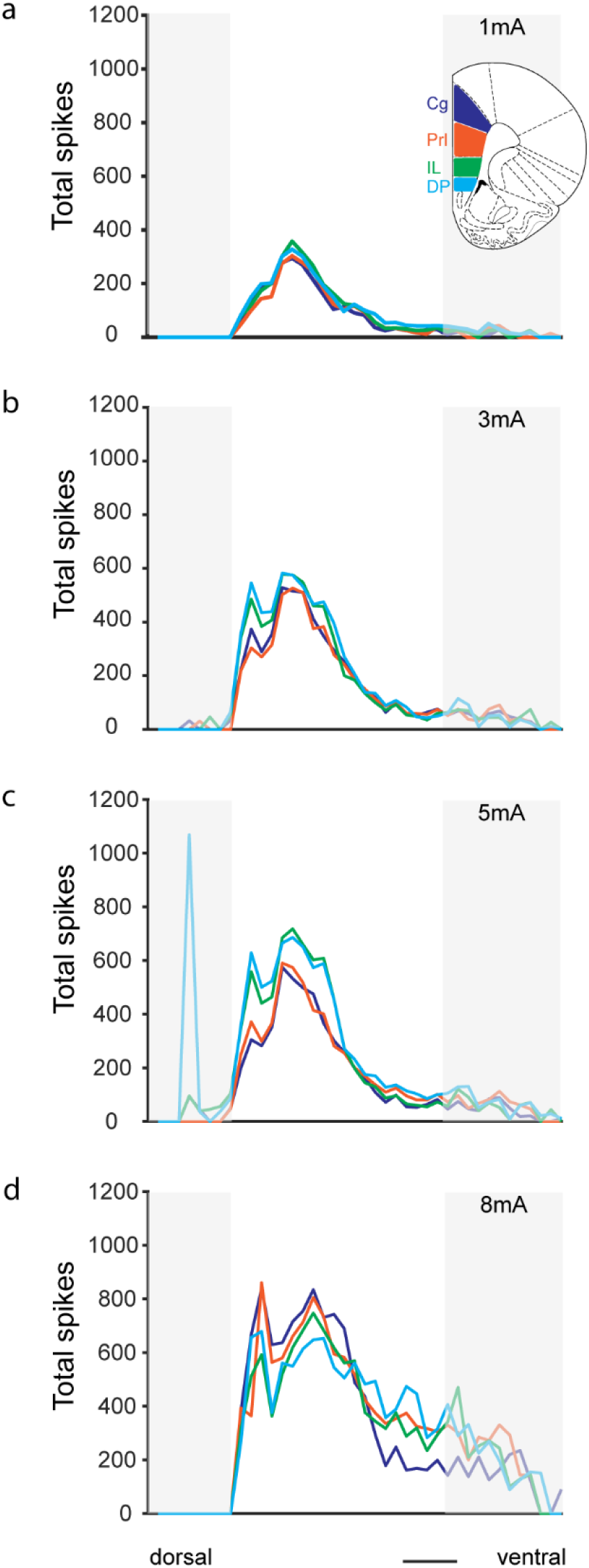
Magnitude of the response in the ICc to electrical stimulation of subregions of the prefrontal cortex at **a** 1mA, **b** 3 mA, **c** 5 mA and **d** 8mA. Subregions are coloured. Note that the same channels are activated from all subregions and that the magnitude of the response is marginally greater in IL and DP than in Prl and Cg. Data are from 8 and 9 stimulation/recording site pairings in 6 animals. Data greyed out are from channels dorsal (left) and ventral (right) to the ICc. Scale bar 500 µm.

### Contralateral cortical regions also have functional connections with ICc

In our previous tracing study, we observed substantial innervation of ICc from the contralateral cortex. Hence, in three animals we examined responses to stimulation at contralateral sites. In two animals we tested contralateral M1, M2 and S1Jaw and found excitatory responses in ICc from all areas. There were short, fixed latency events on some channels and longer latency, longer duration increases in firing rate on many channels. The latencies of the early and later events were not different to those following stimulation of the ipsi-lateral cortex. In two animals we tested contralateral PFC stimulation and found early and later excitatory responses in ICc in one case but not the other. The latencies of these responses were not different from those seen in response to stimulation of the ipsi-lateral PFC.

## Discussion

Our results provide functional evidence that connections from multiple non-auditory cortical areas influence the responses of neurons in the IC. These data are consistent with our previously reported anatomical evidence that the ICc receives descending connections from multiple non-auditory cortical areas (Olthof et al., 2019).

Injection of retrograde virus encoding GFP and ChR2 into the ICc resulted in GFP expression in substantial numbers of pyramidal cells in layer V in AC as well as in multiple non-auditory, cortical regions confirming our previous findings using conventional tracers (Olthof et al., 2019). However, we found that despite injecting virus over a wide area of ICc, cortical labelling was somewhat discrete with small clusters of neuronal somata expressing GFP. In some cortical areas, we could also see GFP in dendrites extending to superficial cortical layers depending on the plane of sectioning. GFP could also be seen in fine axons in the white matter below the cellular layers.

Consistent with the expression of virally encoded ChR2 as well as GFP, blue light stimulation of the cortex evoked firing activity in cortical cells. A ramped increase in light intensity, which minimises desensitization (Adesnik and Scanziani, 2010), evoked substantial increases in firing activity locally in AC, M1, and S1. However, it was of note that, the light-activated regions were discrete, and light failed to increase firing in many cortical regions consistent with the patchy GFP expression. Despite the restricted nature of the cortical activation, optogenetic stimulation of AC and M1 altered firing activity in ICc. The predominant response was an increase in firing rate sometimes evident across several hundred microns. Given that the cortical cells were retrogradely labelled from the ICC and that the virus does not cross synapses (Tervo et al., 2016), this is clear evidence that excitatory inputs to the ICC from AC and from M1 can directly influence the firing of neurones in the ICC. The optogenetic approach arguably provides the best level of evidence however, only small numbers of cortical cells could be activated at any one time, probably because of a combination of restricted light spread within the cortical tissue and patchy viral transfection. Coupled with the likelihood that cortical neurones have a restricted innervation pattern in the ICc, the probability of activating cortical neurones and simultaneously recording from their restricted targets using optogenetics is low. Electrical stimulation avoids some of these issues and allowed us to explore the temporal aspects of the influence of cortical regions on ICc firing, that is not possible with optogenetic stimulation.

Electrical stimulation of multiple non-auditory cortical regions evoked excitatory responses in the ICc. The difference in ‘successful activation’ between electrical stimulation and optogenetic stimulation is likely explained by the greater spread of current and larger area of cortex depolarized by electrical stimulation versus optogenetic stimulation. Nevertheless, there were areas in the cortex from which electrical stimulation failed to evoke responses at the recording sites in ICc. The nature of the responses to stimulation of motor and somatosensory cortical regions was relatively consistent. We frequently saw a singular short and fixed latency firing event of consistent size and shape which occurred with increasing probability as the stimulation current was increased. The short, fixed latency of the event suggest it is monosynaptic (Syka and Popelar, 1984; Qi et al., 2020) and its invariant size suggests that it represents a single spike from a single ICC neurone. The latency of this event decreased with increasing current consistent with higher intensity currents causing more rapid neuronal depolarization. We propose that such events are the result of activation of individual ICc neurones in response excitation by the innervating cortico-collicular connections. Although we did not perform collision tests (Lipski, 1981), it is unlikely that the short latency event is antidromic as there is no direct projection from the ICc to the cerebral cortex (Oliver and Cant, 2018).

Independent of short latency events, the predominant response following electrical stimulation was a longer latency (5-10 ms), longer duration (20-50 ms) increase in firing probability. It was clear from the presence of spikes of different sizes, shapes, and polarity, that this late period of increased firing involved multiple units. We also observed considerable temporal ‘jitter’ in the timing of individual spikes relative to the stimulus confirming that these responses are not antidromic and suggesting that they are polysynaptic in nature. The relatively long latencies observed would also suggest the involvement of multiple neurones activated by synaptic transmission. It is unlikely that the later ‘polysynaptic’ events occur secondary to activation of a cortical circuit since the cortical regions from which the ICc can be activated are discrete. Rather, we propose that direct stimulation of cortico-collicular projections results in short latency monosynaptic activation of individual targeted connections in the ICc which in turn activates local circuits. The increase in response magnitude seen with increasing current likely reflects more consistent monosynaptic activation. With increasing current, excitatory responses also occurred on increasing number of channels particularly in more ventral regions of ICc, albeit the magnitude of responses on the channels recruited with higher currents was always lower than on those channels affected at lower currents. It seems likely that, with higher currents, the greater response magnitude results in further spread of local circuit activation within the ICc such that neurones distant from the direct target of the cortico-collicular neurones become activated.

All our recordings were made in the relative silence of a sound-attenuated room: conditions in which spontaneous activity in the ICc is very low and excitatory responses are most easily captured. However, as has been reported with electrical stimulation of auditory cortex (Syka and Popelar, 1984; Jen et al., 1998) we sometimes saw inhibition of firing associated with electrical stimulation. In the main, this inhibition occurred at longer latencies and was long lasting suggesting a polysynaptic mechanism within ICc. There are large numbers of GABAergic cells within the ICc which modulate the firing of other ICc neurones (Merchán et al., 2005). It is likely that post-stimulus inhibition involves these neurones.

We examined the effects of stimulation of multiple cortical areas on ICc firing. In many cases with the recording electrode in the same position in ICc, we found excitatory responses to stimulation of different motor and somatosensory cortical areas occurred on the same channels. This suggests that the same small regions of ICc, or even the same individual neurones, in ICc are innervated by multiple cortical areas. Consistent with this, we have previously seen terminals anterogradely labelled from different cortical areas, surrounding individual ICc neurones (author’s unpublished observations). It was also notable that stimulation of the different subregions of the PFC evoked responses in the same channels in ICc. Responses were also of similar magnitude and latency suggesting multiple descending cortico-collicular connections from PFC to individual neurons in ICc.

While most of our experiments involved ipsilateral electrical stimulation, we also examined whether ICc neurones would respond to electrical stimulation of contralateral cortical areas. In agreement with our tracing data (Olthof et al., 2019) showing substantial contralateral cortical innervation of ICc, we saw clear excitatory responses to contralateral stimulation which were similar in their nature and timing to those evoked by ipsilateral stimulation.

The ICc is organised tonotopically with more dorsal parts tuned to low frequency sounds and more ventral parts tuned to higher frequencies (Merzenich and Reid, 1974; Malmierca et al., 1995). It was noteworthy that for all the cortical areas studied, greatest responsiveness was observed in low frequency tuned areas of ICc. For M1, M2, and S1FL the largest responses were in channels recording units tuned to around 4 kHz, whereas for simulation of S1Jaw the greatest responses to were in channels recording units tuned around to 1-2 kHz.

The aim of this study was to elicit functional evidence for cortico-collicular connections from non-auditory cortical areas to the IC. As such, our analysis does not demonstrate how these connections influence the responses to sounds in the IC, and thus does not address their physiological significance. As discussed in the Introduction, several studies have reported the modulation of sound responses in IC by non-auditory sensory and motor inputs (Porter et al., 2007; Gao et al., 2015; Lesicko et al., 2016; Leong et al., 2018; Yang et al., 2020; Lohse et al., 2022). The responses we observed may provide the substrate for some of these observations and add to the evidence that influences from non-auditory modalities in the cortex operate at the brainstem level. For example, in the case of the projections from the motor and somatosensory cortices, such circuits may mediate mechanisms that distinguish self-generated sounds from external sources, enhance modality selection, or emphasise signals associated with co-activation of multiple modalities. Furthermore, direct projections from the PFC demonstrate that even regions concerned with executive function can influence midbrain auditory processing.

## Author contributions

SEG, BMJO, and AR designed the research and performed the experiments; SEG and BMJO. analysed the data; SEG wrote the first draft of the paper; SEG, BMJO, and AR edited the paper.

SEG, AR, and BMJO are joint senior authors

## Acknowledgments

This work was supported by the BBSRC (Grant BB/P003249/1 to AR and SEG)

